# Integrated single-cell transcriptomics and chromatin accessibility analysis reveals novel regulators of mammary epithelial cell identity

**DOI:** 10.1101/740746

**Authors:** Nicholas Pervolarakis, Quy H. Nguyen, Guadalupe Gutierrez, Peng Sun, Darisha Jhutty, Grace XY Zheng, Corey M Nemec, Xing Dai, Kazuhide Watanabe, Kai Kessenbrock

**Author notes:** Address correspondence to: Kai Kessenbrock, Ph.D. Kazuhide Watanabe, Ph.D.

## Abstract

The mammary epithelial cell (MEC) system is a bi-layered ductal epithelial network consisting of luminal and basal cells, which is maintained by a lineage of stem and progenitor cell populations. Here, we used integrated single-cell transcriptomics and chromatin accessibility analysis to reconstruct the cell types of the mouse MEC system and their underlying gene regulatory features in an unbiased manner. We define previously unrealized differentiation states within the secretory type of luminal cells, which can be divided into distinct clusters of progenitor and mature secretory cells. By integrating single-cell transcriptomics and chromatin accessibility landscapes, we identified novel cis- and trans-regulatory elements that are differentially activated in the specific epithelial cell types and our newly defined luminal differentiation states. Our work provides an unprecedented resource to reveal novel cis/trans regulatory elements associated with MEC identity and differentiation that will serve as a valuable reference to determine how the chromatin accessibility landscape changes during breast cancer.

## INTRODUCTION

Breast cancer is a heterogeneous disease that can be classified into at least six different intrinsic subtypes, namely luminal A, luminal B, HER2-enriched, basal-like, normal breast and claudin-low (Perou et al., 2000). Breast cancer arises from the breast epithelium, which – in both humans and mice - forms a ductal epithelial network consisting of two main cellular compartments, an inner layer of luminal cells and an outer layer of basal/myoepithelial cells (Visvader, 2009). A series of recent reports have indicated that further heterogeneity exists within these two cell layers in mice. For example, several studies have identified a functionally distinct subpopulation of mammary stem cells within the basal compartment (Shackleton et al., 2006; Stingl et al., 2006). Within the luminal compartment, a subpopulation of progenitor cells has been identified by high expression of *Kit*, and in addition a population of mature luminal cells have been identified using flow cytometry isolation strategies (Shehata et al., 2012).

Advances in next generation sequencing and microfluidic-based handling of cells and reagents now enable us to explore cellular heterogeneity in an unbiased manner using single-cell mRNA sequencing (scRNAseq) to reconstruct transcriptional programs in individual cells (Pollen et al., 2014). Recent studies have utilized this approach to describe the cell types and states within the human (Nguyen et al., 2018) and mouse mammary epithelium (Bach et al., 2017; Pal et al., 2017) generally yielding three main cell types, namely basal (marked by *Krt14*), secretory luminal or also called luminal progenitors (L-Sec; marked by *Elf5*) and mature, hormone-responsive luminal cells (L-HR; marked by *Prlr*). It remains elusive whether additional cellular diversity exists within these three cell types.

In addition to transcriptional programs, cellular identity may be strongly influenced by the epigenetic wiring of the cell, which is not detectable in scRNAseq data. Some of these features may be interrogated systematically by the Assay for Transposase-Accessible Chromatin using sequencing (ATACseq) to reconstruct cis/trans regulatory elements associated with cellular identity (Buenrostro et al., 2015a). Recent advances enabled single cell-level ATACseq (scATACseq) to profile cellular heterogeneity on an epigenetic level (Buenrostro et al., 2015b). This approach enabled unprecedented insights into the differentiation trajectories of the hematopoietic system (Buenrostro et al., 2018; Satpathy et al., 2019), and has recently elucidated transcriptional regulators of developmental lineages of the fetal mammary gland (Chung et al., 2019).

The goal of the present study is to elucidate the molecular underpinnings mediating cellular identity within the mouse mammary epithelium by integrating massively parallel single-cell transcriptomics (scRNAseq) and chromatin accessibility (scATACseq) profiling of isolated mammary epithelial cells (MECs). Our combined single-cell RNA/ATACseq analysis allowed us to identify previously unrealized cell state distribution within the luminal mammary epithelial compartment, and revealed novel cis- and trans-regulatory elements that are associated with cellular identity and differentiation from luminal progenitor into mature, secretory cells. Our work provides novel insights into the spectrum of cellular heterogeneity within the mouse mammary epithelial system under normal homeostasis and will serve as a valuable resource to understand how the system changes during early tumorigenesis and tumor progression.

## RESULTS AND DISCUSSION

### Single-cell chromatin accessibility reveals previously unrealized luminal epithelial cell states in the mouse mammary epithelium

Our recent single-cell transcriptomics analysis of the human MEC system described three distinct cell types of epithelial cells (Nguyen et al., 2018). This pattern is largely conserved in the murine system, as shown in recent scRNAseq analyses of isolated mouse MECs describing three main cell types, namely basal (marked by *Krt14*), secretory luminal (L-sec; marked by *Elf5*) and mature, hormone-responsive luminal cells (L-HR; marked by *Prlr*) (Bach et al., 2017; Pal et al., 2017). Here, we used massively parallel, droplet-enabled scATACseq analysis (10X Genomics Chromium) to determine whether additional cell types and states can be observed on an epigenetic level, and to determine whether cis-regulatory elements and transcription factor motif accessibility are systematically linked with cellular identity in the mammary epithelium. We profiled in total 23,338 individual cells for their chromatin accessibility landscapes. For initial quality validation purposes, we purified basal and luminal MECs by flow cytometry and subjected them to scATACseq analysis in three separate samples each (Figure 1A; Figure S1A-B). After processing the sequencing data using the Cell Ranger pipeline (10X Genomics), we performed unbiased clustering analysis on all peaks using Seurat, which revealed 4 main clusters (0-3) of MECs (Figure 1B), as well as minor populations of contaminating stromal cells that were excluded from this visualization. To identify cell types, we generated a gene activity matrix to serve as pseudo-expression data as previously described (Stuart et al., 2019). This enabled us to identify basal cells (cluster 0; marked by *Krt14*), L-Sec (Clusters 2-3; marked by *Kit*) and L-HR (cluster 1; marked by *FoxA1*) (Figure 1C; Figure S1C).

**Figure 1:**
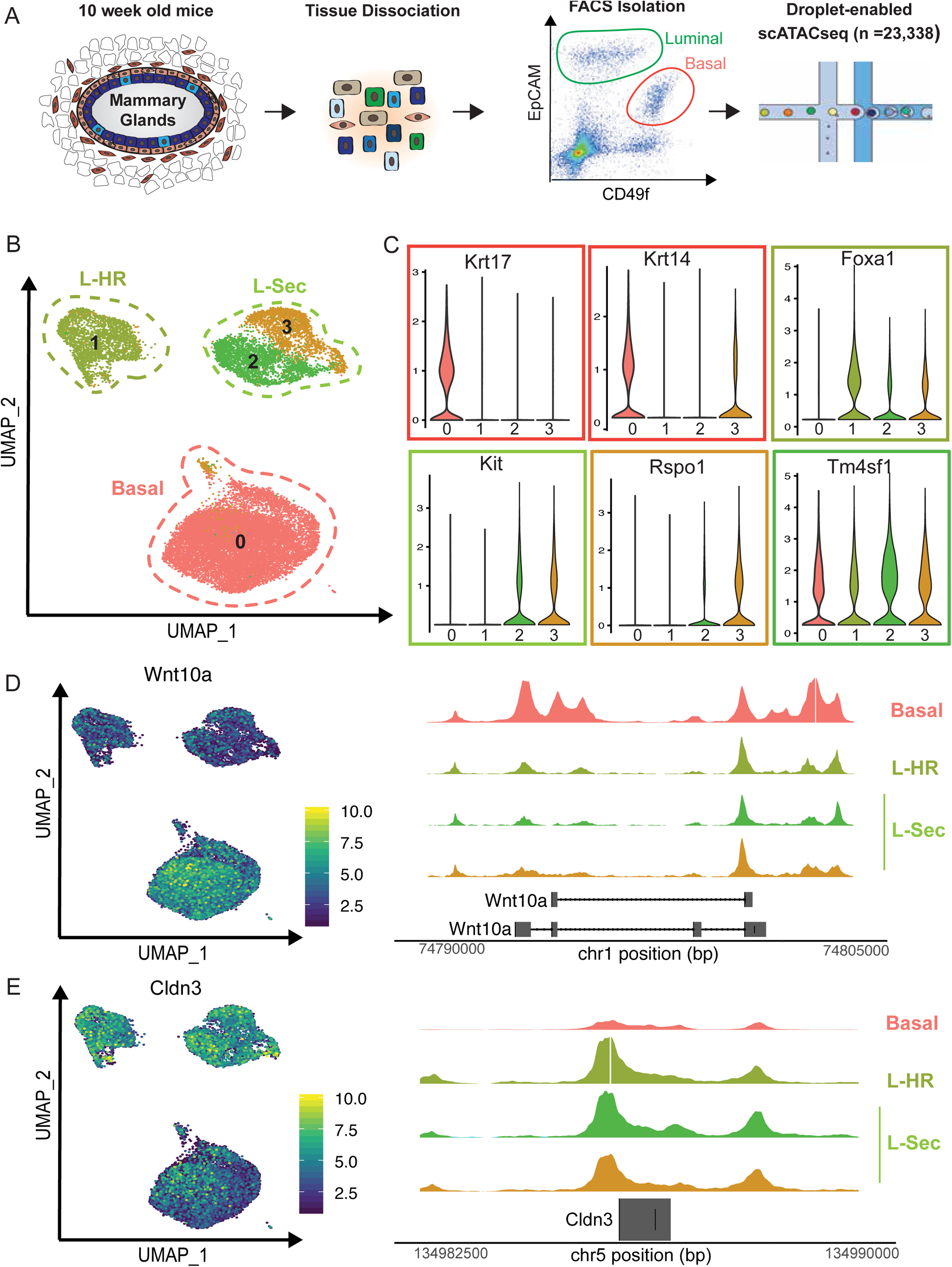
(A) Schematic of experimental workflow for scATACseq analysis. (B) UMAP visualization of scATACseq libraries, colored by Seurat clustering performed on aggregated peak matrix. Cell types are outlined in dotted lines with Basal in red, Luminal-Hormone Responsive (L-HR) in light green, and Luminal-Secretory (L-Sec) outlined in dark green. (C) Violin plots of Cicero-generated gene accessibility matrix-based marker genes of each cluster, with boxes colored by cell type-specific accessibility. (D-E) UMAP of scATACseq analysis on the left, with cells colored by gene accessibility expression level of Wnt10a and Cldn3. Pseudo-bulk profiles of library fragments on the right, subset by cluster at genomic regions corresponding to Wnt10a and Cldn3.

Interestingly, we observed two previously unrealized distinct clusters within the L-Sec cell type, which contained cluster 2 marked by *Tm4sf1* encoding a tetraspanin transmembrane molecule involved in breast cancer metastasis through regulation of PI3K pathway (Sun et al., 2015), while cluster 3 displays accessibility of the gene *Rspo1* encoding the regulator of Wnt signaling R-Spondin 1 that plays an important role in mediating mammary stem cell renewal (Cai et al., 2014). Further gene activity analysis enabled us to define numerous specific marker loci that are specifically accessible in the clusters (Figure S1C). Interestingly, cluster 3 also showed moderate accessibility of the basal marker gene *Krt14* (Figure 1C), suggesting that - although of luminal epithelial nature according to flow cytometry and other hallmark gene activity such as *Krt8/18* - this cell state within L-Sec shows some similarity to basal cells. This was particularly intriguing to us, since it could indicate a bipotent progenitor cell state that possesses the capacity to differentiate into both basal and luminal lineages, or a transitory luminal progenitor cell type that is directly derived from a basal mammary stem cell as previously proposed (Shackleton et al., 2006; Stingl et al., 2006).

Finally, we generated pseudo-bulk profiles to visualize genomic regions that were differentially accessible between basal and luminal MECs to reveal the substructure of chromatin accessibility variation between the two main basal and luminal compartments. Focusing on *Wnt10a*, which was found to be specifically open in basal cells based on our gene activity analysis (Figure 1D), we found that the main difference between basal and luminal MECs was observed around the immediate gene promoter region, while there was no difference observed around the terminal region of *Wnt10a*. Similarly, the luminal-restricted gene *Cldn3* displayed one major peak of high accessibility in proximity to the gene promoter in all three clusters of luminal cells, which was essentially absent in the basal pseudo-bulk analysis (Figure 1E). Taken together, these initial analyses showed that our scATACseq dataset represents a valuable resource to explore the chromatin accessibility landscape in individual mouse mammary epithelial cells. Using our dataset, we define two previously unrealized distinct cell states characterized by differential chromatin accessibility features (e.g. *Rspo1*) within the secretory luminal epithelial cell type (L-Sec).

### Defining the distinct gene expression signatures within MEC cell types and states using single-cell RNA sequencing

To further explore the distinct gene expression signatures underlying the cell types and states revealed by scATACseq, we performed scRNAseq on FACS-isolated MECs from age- and background-matched 10-week old, female FVB/NJ mice yielding a dataset of 26,859 single-cell transcriptome libraries (Figure 2A; Figure S2A-B). Using unbiased Seurat clustering, we detected three main clusters of epithelial cells (Figure 2B) that correspond to basal (*Krt14*+), secretory luminal (L-Sec; *Kit/Elf5*+) and luminal, hormone-responsive cells (L-HR; *Prlr*+), which is in line with previous single-cell transcriptomics analyses of mouse MECs (Bach et al., 2017; Pal et al., 2017). All clusters were evenly composed of cells from all three individual experiments, which confirms that the cell states defined here are highly reproducible (Figure S2C). We also detected a small cluster of contaminating stromal cells expressing various non-epithelial genes, and minor clusters of proliferating cells mainly between L-Sec and basal clusters (*Mki67*+; Figure S2C). In addition, we noticed small clusters of cells expressing both luminal and basal keratins and displayed high levels of genes per cells suggesting that these represent doublets (D). Interestingly, we detected two distinct cell states within the L-Sec cluster of MECs (Figure 2B), which emerged as one homogeneous cluster in previous scRNAseq studies of mouse MECs (Bach et al., 2017; Pal et al., 2017). Further interrogation of specific marker gene expression (Figure S2C; **Table S1**) revealed that one of these clusters expressed several genes associated with milk production (*Lipa*, *Csn2*, *Lalba*), while the second cluster expressed high levels of genes associated with epithelial progenitor cell capacity (*Aldh1a3*, *Rspo1*). We therefore named these sub-clusters “L-sec Progenitor” and “L-sec Mature” (Figure 2B-C). These designations were further supported by the top Gene Ontology (GO) terms associated with their marker gene signatures, namely “secretory granule (GO: 0034774)” and “ribosome (GO: 0005840)”, which is in line with observation that progenitor cell populations are characterized by increased ribosomal gene expression in the absence of cell proliferation (Athanasiadis et al., 2017). Since aldehyde dehydrogenase expression was previously shown to mark a subset of luminal-restricted progenitor cells in the human breast (Eirew et al., 2012), we next used Aldh1a3 as a marker for *in situ* validation of this cell population. Using a specific RNA-based probe (RNAscope) for *Aldh1a3* in combination with anti-KRT14 antibody staining to label the basal cell compartment, we detected a subset of luminal epithelial cells (KRT14-negative) with pronounced expression of *Aldh1a3* located in both ductal and lobular regions of the mammary gland (Figure 2D). Quantification of cells and numbers of *Aldh1a3* transcripts detected by RNAscope revealed a frequency of high-expressing cells that was very similar to our scRNAseq results (Figure 2E). Taken together, these findings confirm the existence of two distinct states within the L-sec cell type as predicted by scATACseq, and allowed us to integrate these results with previously proposed functional designation as luminal progenitor and mature secretory luminal cells.

**Figure 2:**
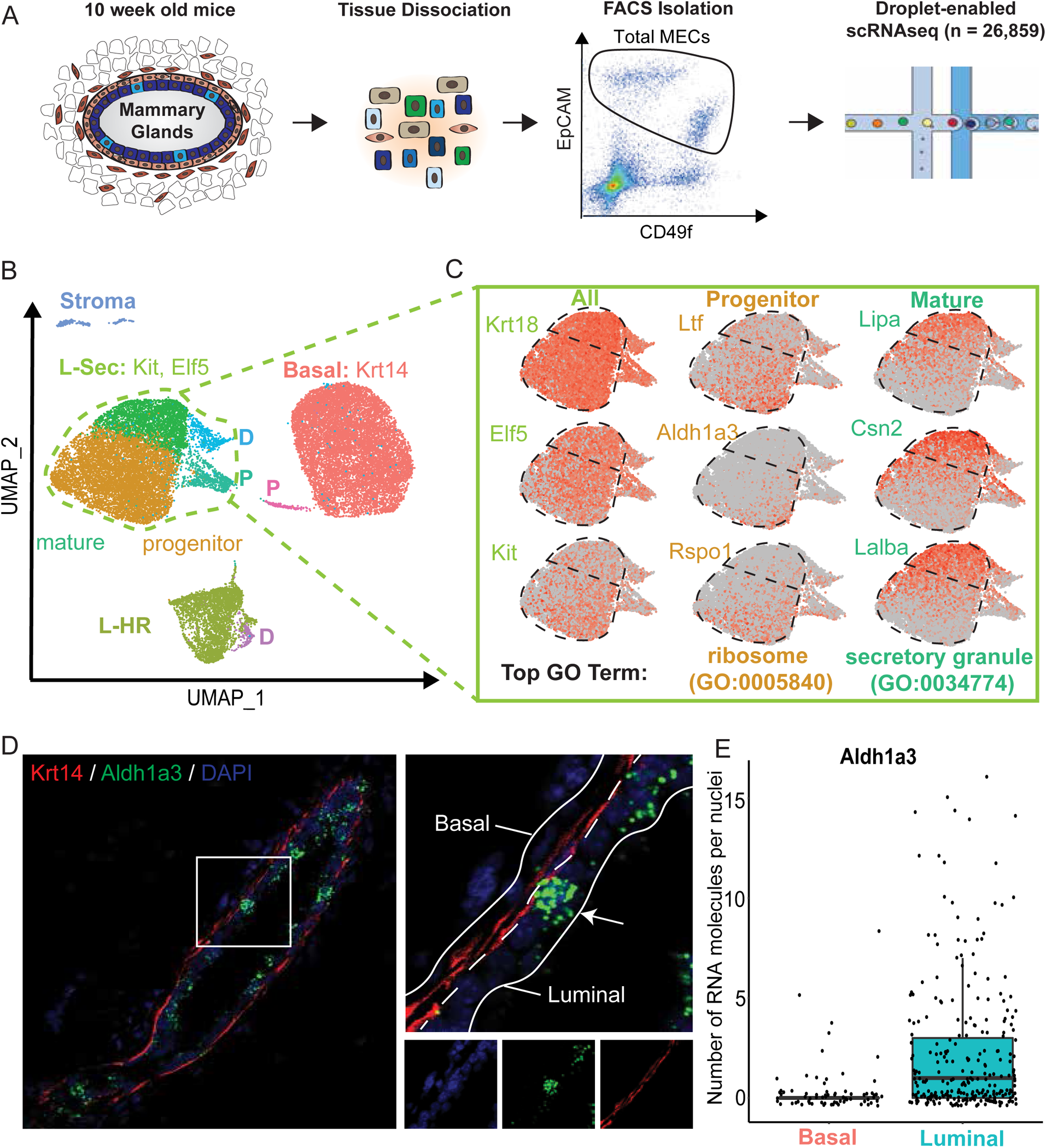
(A) Schematic of experimental workflow for scRNAseq analysis of isolated mouse MECs. (B) UMAP visualization of scRNAseq libraries anchored by sample, with colors corresponding to unbiased clustering and annotated by cell type and state. Basal cells are in red, L-HR in light green, and L-Sec outlined in dark green. Putative doublets are marked by the letter D, and proliferative cells are marked by the letter P. Within L-Sec cell type, two distinct clusters emerged that were labeled as Mature and Progenitor based on gene expression signatures. (C) Focused analysis of L-Sec cluster and corresponding marker gene is shown, where expression is scaled such that dark red corresponds to high expression of the gene and light grey corresponds low expression of the gene in question; top GO-term including accession number is listed for Progenitor and Mature cell states. (D-E) Fluorescence images from *in situ* RNAscope analysis for *Aldh1a3* in combination with immunostaining for basal-specific KRT14 is shown and luminal and basal compartments are outlined in blown up image. Quantification of transcript counts per basal and luminal cells is shown; data was combined from three independent ductal regions of mouse mammary gland sections.

### Integration of single-cell RNA and ATAC sequencing reveals novel cell type-specific transcriptional regulators and cis-regulatory elements

We next sought to integrate our scRNAseq and scATACseq datasets to gain deeper biological understanding about the link between chromatin accessibility and gene expression within mammary epithelial cell types. To this end, we utilized a previously described approach to “anchor” diverse datasets together for comprehensive integration of single-cell modalities (Stuart et al., 2019). In short, transfer anchors were learned between the scRNAseq dataset and scATACseq-based gene activity matrix, using scRNAseq as the reference. Cluster labels from scRNAseq were predicted using these anchors in the scATACseq cells yielding a high degree of overlap between expected labels based on markers and the previously assigned cluster labeling (Figure S3A). Next, the integration anchors were used to generate an imputed expression matrix for the scATACseq cells, and the resulting data was merged with our scRNAseq analysis. Visualization of this integrated dataset yielded consistent overlap between both modalities within each of the main cell types, and nicely recapitulated the two clusters of progenitor and mature cells within the L-Sec cell type (Figure 3A, Figure S3B). We observed overall high correlation between ATACseq and RNAseq data in each of the defined cell types (Figure S3C). We next explored several known hallmark genes for cell types in the mammary gland (e.g. *Krt5*, *Krt8*, *Kit*, *Foxa1*) using this integrated analysis, which showed strong correspondence between chromatin accessibility and gene expression in each cell type (Figure 3B). To further corroborate the progenitor and mature cell state within L-Sec, we observed striking consistency for *Rspo1* in progenitor cells and *Lalba* in mature L-Sec cells in terms of chromatin accessibility paired with gene expression (Figure 3C).

**Figure 3:**
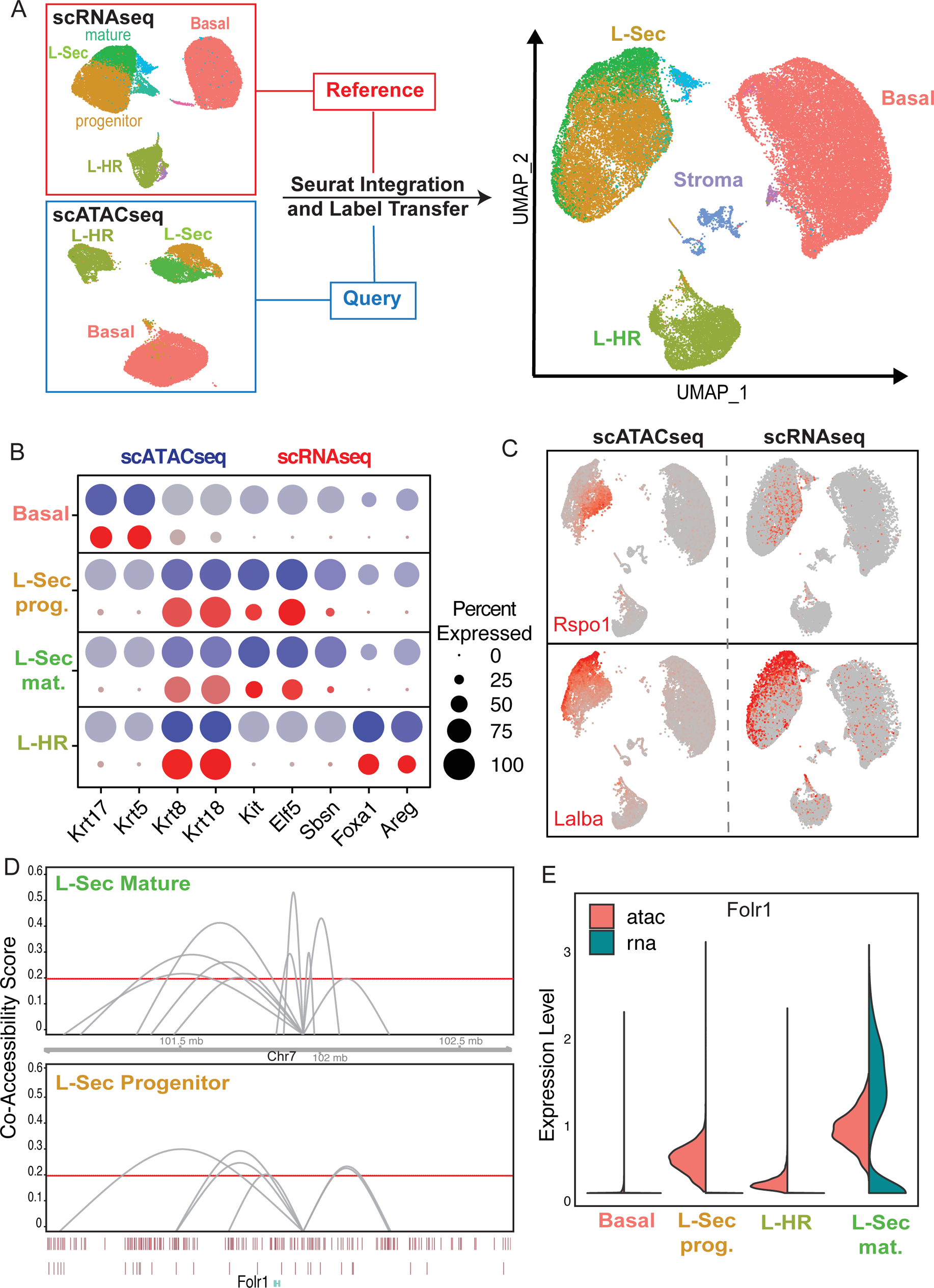
(A) Schematic of Seurat-based label transfer of scRNAseq cell clusters onto scATACseq cells, and subsequent coembedding into a single UMAP visualization. (B) Split Dot Plot of cell type markers, with imputed RNA expression intensity in the scATACseq cells scaled from grey to dark blue, and expression intensity in scRNAseq cells displayed in a scale from grey to dark red. The size of the dot corresponds to the percentage of cells within that cluster have positive expression of the gene. (C) Coembedded UMAP faceted by library type, with cells from scATACseq libraries on the left and cells from scRNAseq libraries on the right. Cells are colored based on scaled expression, with grey corresponding to low and dark red corresponding to high expression. (D) Cicero connection data at enhancer region chr7_101932449_101936345 generated by subset analysis by cluster. Connections from L-sec Mature cells shown in top panel, and connections from L-sec Progenitor shown in bottom, with a minimum co-accessibility score of 0.2 visualized. (E) Violin plot of *Folr1* expression in the coembedded analysis, split by technology type.

To identify cis-regulatory elements that may contribute to cell type distinction, we used the Cicero pipeline for co-accessibility analysis to determine cell type-specific genomic connections (Pliner et al., 2018). This analysis generates a data frame of pairs of peak regions and calculates a score of how frequently these are both accessible in the same cells. The resulting connections were subset by those in which one peak of each pair corresponded to an enhancer region from EnhancerDB’s mouse mammary putative enhancer list (Gao et al., 2016). Connections above a co-accessibility score of 0.5 were selected, and the closest protein coding regions of the non-enhancer genomic region were annotated. Directly comparing L-Sec Mature and L-Sec Progenitor cells, we found enhancer-specific connections near the *Folr1* locus that were specific to the L-Sec Mature population, but not the L-Sec progenitors (Figure 3D). Further interrogation of gene expression and chromatin accessibility revealed specific signal for *Folr1* in L-Sec mature (Figure 3E). Interestingly, *Folr1* was recently identified as a putative regulator of milk protein synthesis in cow mammary glands (Menzies et al., 2009), which is in line with the high degree of secretory and lactation-associated genes in the L-Sec mature cells (Figure 2). Together, this suggests that this enhancer region on Chromosome 7 represents a key regulatory element that becomes active during differentiation into mature secretory luminal MECs. In addition, our approach also revealed a basal-specific enhancer region connected with the *Cnn2* gene locus that had no connections present in any of the other cell types, and showed basal restricted expression of *Cnn2* (Figure S3D-E). Since calponin isoforms CNN1, CNN2 and CNN3 regulate cytoskeleton functions in smooth muscle cells (Liu and Jin, 2016), this may be a critical feature for mediating the myoepithelial nature associated with basal MECs. Future experiments will pursue functional evaluation of the significance of these enhancer regions for mammary epithelial differentiation and cell type identity.

We next sought to identify transcription factors (TFs) that may be critical for regulating mammary epithelial cell type identity. We utilized the ChromVar analysis pipeline (Schep et al., 2017) to analyze accessibility of TF motifs in our scATACseq dataset that are specific for each cell type (**Table S2**). Trimming the output to those TF’s that were significantly associated with a cell type through logistic regression, we performed co-correlation analysis to pinpoint down TF modules in the MEC system (Figure 4A). This approach revealed three major modules: Module 1 contained predominantly Jun and Fos-related TF motifs indicating that this feature is related to a subset of cells showing stress-response most likely due to tissue dissociation and FACS isolation; Module 2 contained numerous TFs previously associated with basal epithelial biology such as Tp63 (Forster et al., 2014), but in addition Gata3 and other Gata family TFs were observed, which have been linked with regulating luminal cell fate decisions (Kouros-Mehr et al., 2006); finally Module 3 contained mostly TFs associated with luminal epithelial biology such as Foxa1 (Liu et al., 2016) and Elf5 (Zhou et al., 2005), but also included a cluster of EMT-related TFs such as Tcf4, Snai2 and ID4 (Stemmler et al., 2019).

**Figure 4:**
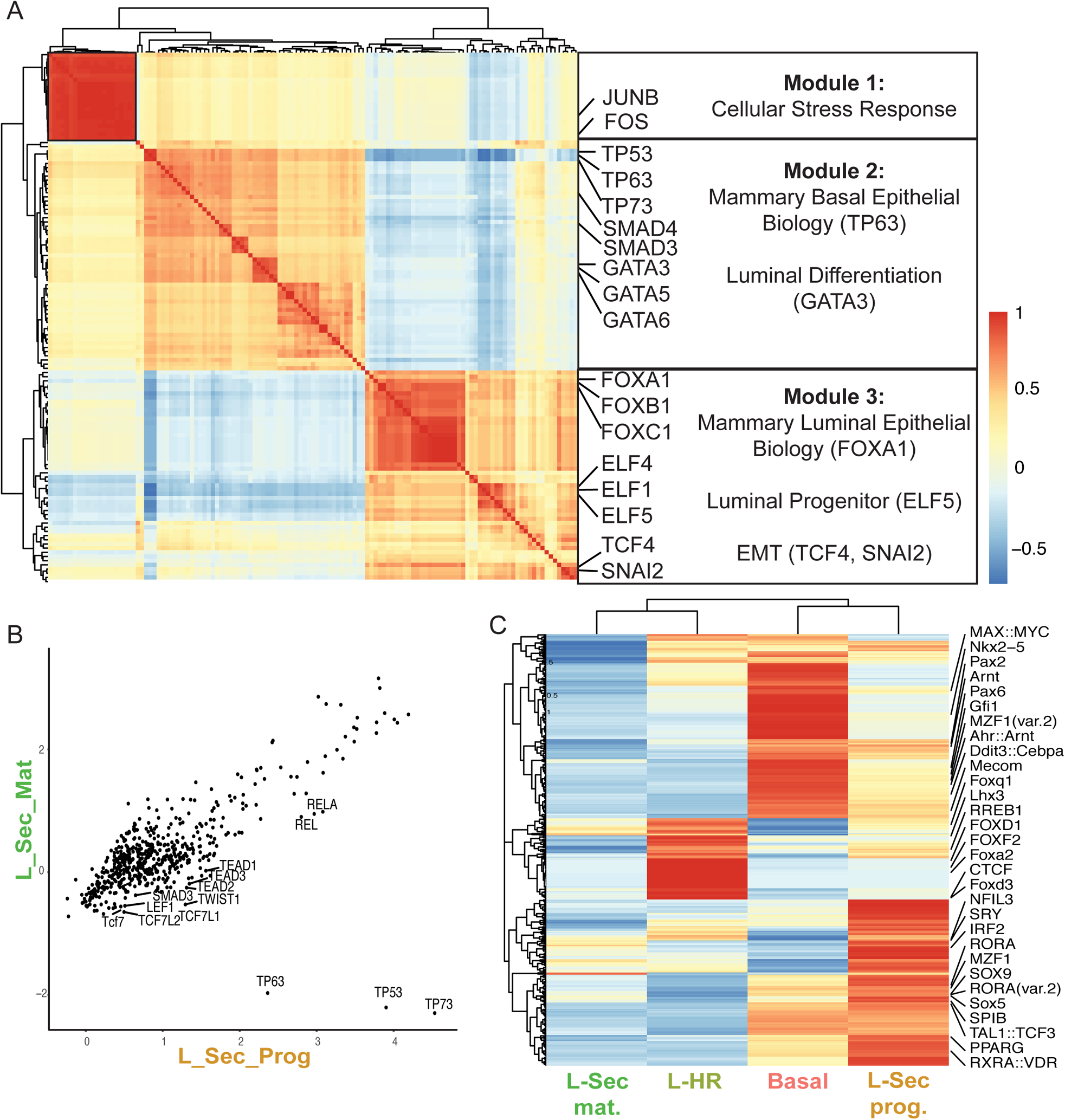
(A) Heatmap displaying co-correlation of TF motif accessibility shown as z-scores (blue = low; red = high). Transcription factors were subset to those that were found through logistic regression to be significantly associated with a particular cluster post label transfer from scRNAseq data onto scATACseq data, and had an average log-fc greater than one. Key TFs are highlighted in relation to putative function on the box to the right. (B) Comparison of TF Motif z-scores between L_Sec_Mat vs. L_Sec_Prog cell populations. Selected TFs are labeled. (C) Hierarchical clustering of cell type-specific TF Motif z-scores is shown; blue = low and red = high.

To devise TFs that may play a central role in regulating MEC identity, we next focused on TFs that within the same cell type display both motif accessibility as well as high scores for an active downstream target gene expression as determined by Enrichr analysis (**Table S3**). Very reassuringly, the master regulator of basal/myoepithelial cell biology Tp63 (Forster et al., 2014) emerged as one of the top TF motifs that was specifically accessible in basal cells, and also showed distinct gene expression as calculated using gene score for a set of target genes (**Figure S4**). Several SMAD TFs yielded top motif scores within basal cells, however, SMAD3 showed highest target gene expression scores in basal cells indicating that SMAD3 represents a key TF in the regulation of basal cell identity. SMAD family TFs are critical mediators of transforming growth factor β1 (TGF-β), which has wide implications in regulating mammary epithelial biology and cancer (Moses and Barcellos-Hoff, 2011). Our findings place particular importance of SMAD TFs in the context of basal MEC biology. Interestingly, the luminal progenitor-associated TF ELF5 (Zhou et al., 2005) showed highest motif accessibility in both luminal clusters (L-Sec and L-HR), however, expression of ELF5 target genes is most predominantly detected in L-Sec cells. This can be explained by the fact that *Elf5* gene expression is almost exclusively found in the L-Sec cluster (Figure 2C) indicating that while the ELF5 TF motif is accessible in L-HR cells, its downstream gene program is not activated unless the TF itself is expressed. Finally, we explored FOXA1 as a known regulator of luminal differentiation, which indeed showed strong correspondence between high TF motif accessibility and elevated target gene expression scores specifically in L-HR cells corroborating the notion that FOXA1 is a master regulator of the hormone-responsive luminal cell type (Bernardo et al., 2010).

To define the differences between the newly established L-Sec Mature and Progenitor cell states, we specifically compared these clusters for differential TF motif accessibility (Figure 4B), which yielded numerous Wnt signaling-related TFs such as LEF1 and TCF7L2 upregulated in L-Sec Progenitors. This is in line with increased expression of the potentiator of canonical Wnt signaling *Rspo1* in this cluster. In addition, we noticed accessibility for several basal cell-specific TFs such as TP63, which is supported by our observation of increased accessibility of the hallmark basal keratin *Krt14* in this cluster (Figure 1C). To systematically explore the relationship of the MEC cell types and states, we next performed hierarchical clustering based on TF motif accessibility showing that L-Sec Progenitors cluster closely with Basal cells and share many TF modules between each other, while L-Sec Mature more closely align with L-HR cells. These shared features between basal cells and luminal progenitors support the concept that luminal progenitors may be directly derived from basal mammary stem cells that possess the capacity to differentiate into luminal cells (Visvader, 2009). This will need to be further corroborated using functional or targeted lineage tracing experiments.

Taken together, our integrated single-cell transcriptomics and chromatin accessibility analysis of the MEC system revealed a previously unrealized cell type hierarchy within the luminal epithelial compartment, and defined novel transcriptional and epigenetic underpinnings regulating cellular identity in the mammary epithelium. In particular, we define distinct maturation states within the secretory type of luminal cells (L-Sec), which can be divided into progenitor (*Rspo1*, *Aldh1a3*) and mature secretory cells (*Lalba*, *Csn2*). By directly integrating transcriptomics and chromatin accessibility datasets, we were able to provide a framework to devise putative key transcription factors by combining motif accessibility with positive downstream target gene expression. We also identified novel enhancer regions that are systematically associated with gene accessibility and expression of effector genes associated with secretory luminal maturation (*Folr1*) as well as with basal, myoepithelial cell identity (*Cnn2*). Our findings lay the groundwork for future studies to functionally address the biological significance of these cis/trans regulatory elements in mediating mammary stem and progenitor cell function, and to determine how the chromatin accessibility landscape changes during breast cancer.

## METHODS

### Cell Isolation and single-cell RNA and ATAC sequencing library generation

<u>Mice:</u> FVB/NJ mice are from Jackson Laboratory (Stock Number: 001800). In both scRNAseq and scATACseq experiment, 10 weeks old female mice were used for tissues collection. All experiments have been approved and abide by regulatory guidelines of the International Animal Care and Use Committee (IACUC) of the University of California, Irvine.

<u>scRNAseq/scATACseq:</u> Mammary glands number 4 were collected and pooled from a total of four 10-week old, female FVB/NJ mice. Glands were minced into pieces ∼1mm in diameter and processed as previously described (Kessenbrock et al., 2013). In brief, minced glands were incubated with a 2mg/ml collagenase type IV solution at 37C while shaking for 1 hour. Digested organoids were collected by differential centrifugation. Collected organoids were further dissociated with trypsin into single cells. Cells were stained for flow cytometry using fluorescently labeled antibodies for CD49f, EpCAM, CD31, CD45, Ter119, and SytoxBlue. For scRNAseq, live epithelial cells were collected for sequencing. For scATACseq, basal and luminal cells were collected separately.

Library generation for 10x Genomics v2 chemistry was performed following the Chromium Single Cell 3’ Reagents Kits v2 User Guide: CG00052 Rev B. Library generation for single cell ATACseq were performed following the Chromium Single Cell ATAC Reagent Kits User Guide: CG000168 Rev B. Single cell RNAseq and ATACseq libraries were sequenced on the Illumina HiSeq4000 platform targeting approximately 50,000 reads per cells.

### Sequence alignment and data processing

Alignment of scRNAseq analyses was completed utilizing 10x Genomics Cell Ranger pipeline (version 2.1.0). Alignment of scATACseq analyses was completed utilizing 10x Genomics Cell Ranger ATAC pipeline (version 1.1.0). Each library was aligned to an indexed mm10 genome using Cell Ranger Count and Cell Ranger ATAC Count. “Cell Ranger Aggr” function was used to normalize the number of confidently mapped reads per cells across the libraries from different libraries for scRNAseq and scATACseq separately.

### Cell-type clustering analysis and marker identification using Seurat

The aggregated peak-by-cell data matrix was read into R (R version 3.6.0) and processed using the Seurat single cell analysis package version 3.0.2 (Macosko et al., 2015). Along with the peak matrix, the Cicero-generated gene activity matrix (see below) and ChromVar deviations score matrix (see below) were added as assays to the Seurat object. A quality control cutoff of a minimum of 2500 fragments per cell was applied to trim the data set of low-quality cells. Next, variable features of the peak matrix were set to peak regions of >100 across the matrix. These variable features were used to perform Latent Semantic Indexing (LSI), and the first 50 components were calculated. These components were then used to generate a Uniform Manifold Approximation and Projection (UMAP) dimensionality reduction. Post UMAP, a Shared-Nearest-Neighbor graph was generated from the first 14 LSI components chosen via the elbow plot method and was used to cluster the cells via Seurat’s Louvain algorithm.

Marker genes for peak-based clustering were generated using Seurat’s default FindAllMarkers() function on the gene activity matrix. Pseudobulk profiles by cluster highlighting fragment stack ups at particular genomic regions were generated using Signac (version 0.1.0).

Post label transfer, cell type-specific transcription factor motifs were calculated using the logistic regression method option implemented in Seurat’s FindAllMarkers() function. Those TF motifs that had an average log fold change greater than one were used to generate the correlation heatmap to find co-correlated modules of transcription factor motif enrichment.

### scRNAseq Analysis

Each of the scRNAseq data libraries were independently read into R version 3.6.0 and processed using the Seurat pipeline version 3.0.2. Genes had to be expressed in at least three cells to be considered for analysis. Cells were trimmed to those that had at least 200 minimum unique genes expressed, no more than 6000 unique genes, and less than 30% of counts aligning to the mitochondrial genome. Libraries were anchored and integrated using the top 2000 variable features per library calculated via the “vst” method in Seurat. CCA on these 2000 features between the libraries was calculated, and the first 20 dimensions used as input for anchoring. Post anchoring, PCA was performed and the first 10 PC’s were used for UMAP dimensionality reduction and subsequent clustering using the default Louvain implementation. Marker genes per cluster were calculated using Seurat’s FindAllMarkers() function and the “wilcox” test option. GO term enrichment was performed using Enrichr (Kuleshov et al., 2016).

### Gene activity matrix generation

The aggregated peak-by-cell data matrix was read into R version 3.6.0, binarized, and processed with the Cicero analysis package version 1.2.0 and the monocle 3 alpha version 2.99.3 to generate a gene activity matrix for all cells sequenced in the study. The generation of the matrix took into account not only fragments that aligned to regions proximal to the promoter site of each protein coding gene in the genome took into account peak co-accessibility scores also generated through Cicero for all cells to factor in distal genomic relationships to the promoter site of each gene.

### Cis-regulatory regions by cluster

Post label transfer, scATACseq cell libraries were subset by their predicted ID label, whereupon the Cicero pipeline was utilized on each subset. Co-accessibility networks were generated, with pairs of peak regions and their corresponding score in a data frame. This data frame was subset to only those pairs that overlapped with regions in the Enhancer Atlas mouse mammary list as the first peak of the connection (Gao et al., 2016). This trimmed connection matrix was then thresholded for each cell type to those that had a co-accessibility score greater than 0.5. Next, the second non-enhancer peak in the pair was annotated to its closest protein coding gene. Conserved expression markers between technology (RNA and ATAC in the RNA-imputed matrix) were found by cell type and the respective co-accessible gene regions that were both highly connected to an enhancer region, and represented a marker for a cell type were selected.

### Transcription factor (TF) motif analysis using ChromVar

Motif enrichment analysis was performed using an R package ChromVAR version 1.4.1 (Schep et al., 2017). Open chromatin peaks and read counts at open chromatin were defined by the Cell Ranger pipeline as described above. After correction of GC bias, TF deviation score was calculated using a total of 579 TF motif position weight matrices provided with the 10X Genomics Cell Ranger package. For TF clustering analysis, only cells corresponding to epithelial clusters post label transfer (0,1,2,3) were selected. TF enrichment scores were averaged by cluster and hierarchically clustered using hclust() and pheatmap() in R.

### Combined scATACseq and scRNAseq analysis

To generate a coembedding of cells from both scATACseq and scRNAseq libraries, cells from the scRNAseq analysis were used as a reference dataset to predict cluster labels in the scATACseq dataset and transfer them. This prediction used the variable features of the scRNAseq analysis on the RNA assay, and the gene activity matrix of the scATACseq analysis as the query data. Transfer anchors were learned using FindTransferAnchors() and the cluster labels were predicted using the TransferData() function together with the peak-based scATACseq LSI reduction as the weight.reduction function option input. Next, an imputed gene activity matrix was generated by using the TransferData() function again, with the previously learned transfer anchors and a matrix consisting of only the variable features of the scRNAseq analysis and its corresponding cells as the reference. This imputed expression matrix was then used to merge the two Seurat objects, allowing for co-visualization of cells labeled by the scRNAseq cluster labels or their predicted cluster labels for the scRNAseq based or scATACseq respectively.

For combined TF motif accessibility and target gene expression analysis, we first identified cell type-specific TF motifs in our ChromVar analysis (see above), and then performed Enrichr analysis using cell type marker genes from scRNAseq to identify “ENCODE and ChEA Consensus TFs from ChIP-X” for each cell type. Transcription factor targets came from the Enrichr analysis (**Table S3**), where marker genes by cluster were analyzed and those genes that had pathway hits in the Encode database annotation for particular transcription factors were used to score all cells using Seurat’s AddModuleScore() function.

### In situ RNA analysis using RNAscope

Mammary glands were harvested from a 10-week old C57BL/6 mouse and frozen in O.C.T Compound (4583, Sakura). 10-micron sections were fixed with fresh 4% PFA made from 40% PFA (15715-S, Electron Microscopy Sciences) diluted in PBS (21-031-CV, Corning) for 1 hour at RT. The RNAscope assay for the Aldh1a3 probe (501201, ACDBio) was performed according to the manufacturer’s protocol for fresh frozen sections. The images were acquired with a Zeiss LSM 700 confocal microscope. Fiji was used to calculate the number of Aldh1a3 foci (RNA molecules) per nuclei manually. Nuclei enveloped in Krt14 protein are called basal for this analysis. Nuclei adjacent to, but not enveloped, are called luminal.

## Acknowledgements

We thank Devon Lawson for carefully reading the manuscript. This study was supported by funds from the NIH/NCI (1R01CA234496; 4R00CA181490 to K.K.), the American Cancer Society (132551-RSG-18-194-01-DDC to K.K.), and the NIH/NIGMS (R01GM123731 to X.D.). Grant-in-Aid for Scientific Research (KAKENHI) on Innovative Areas “Cellular Diversity” (JP18H05106 to K.W.).

## Author Contributions

K.K. and K.W. designed research and supervised research; Q.H.N., K.W., G.G., P.S., D.J., performed research; X.D., J.R., G.X.Y.Z. and C.M.N. contributed new reagents and analytic tools; N.P., Q.H.N. and K.W. performed bioinformatic analyses; N.P., K.W., and K.K. wrote the paper manuscript, and all authors discussed the results and provided comments and feedback.

## Data Availability

All data will be accessible at Gene Expression Omnibus (GEO accession number #PLACEHOLDER), including raw .fastq files and quantified data matrices along with their associated meta data.

Code Availability: All code will be made available upon request.

---

**Figure S1:**
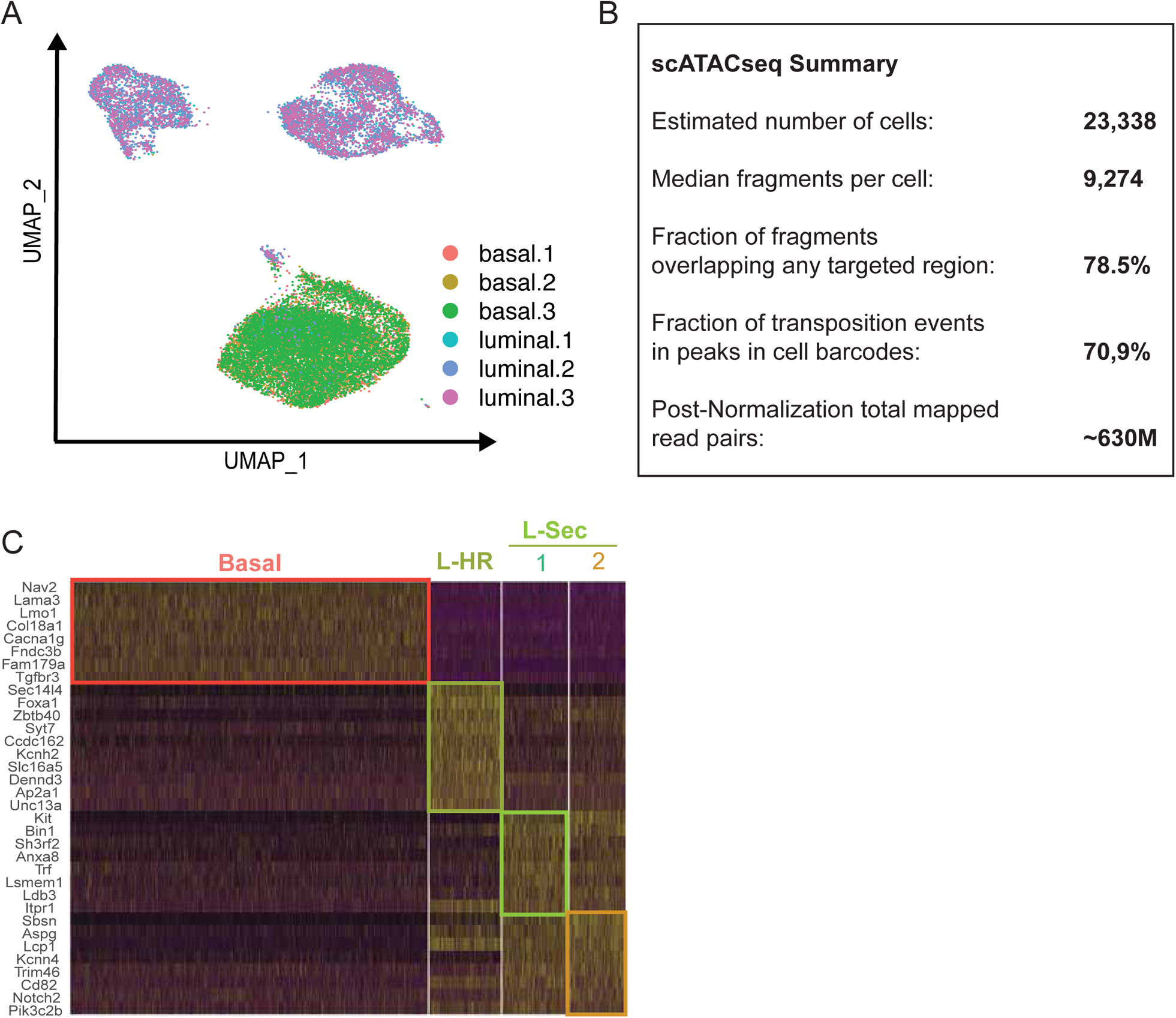
(A) UMAP of scATACseq cell libraries, with cells colored by library of origin. (B) Sequencing and alignment statistics for the six scATACseq libraries. (C) Gene Accessibility matrix top marker gene expression heatmap, with yellow corresponding to high expression and purple corresponding to low.

**Figure S2:**
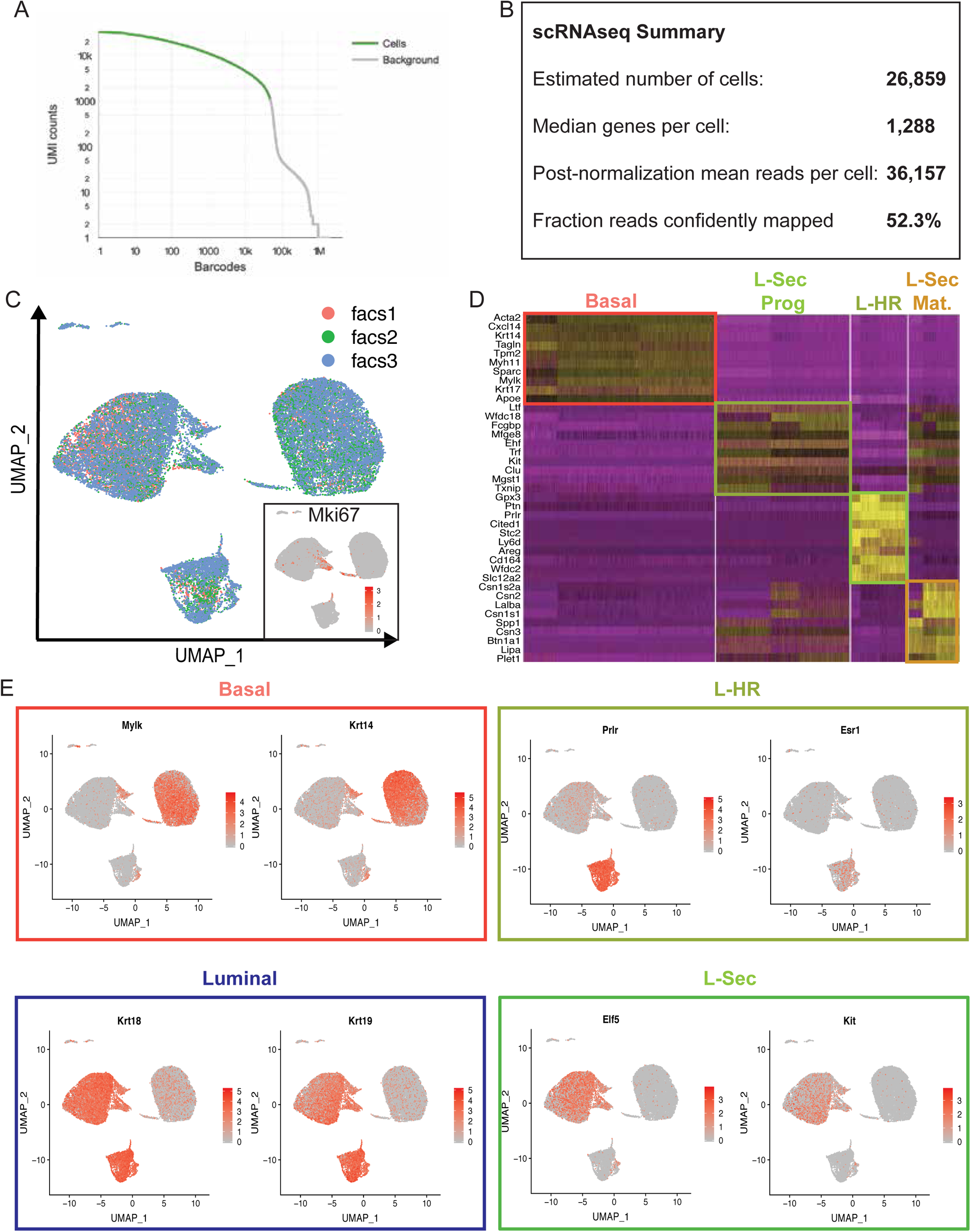
(A) Cell Ranger UMI-based cutoff to determine cell-associated barcodes. (B) Sequencing and alignment statistics for the three scRNAseq libraries. (C) UMAP of scRNA-seq cells, with cells colored by library of origin. Proliferative cells are marked by high *Mki67* gene expression, with dark red corresponding to high expression and light grey to low. (D) Heatmap showing top 10 marker genes per cluster (yellow = high expression and purple = low expression). (E) Feature plots in UMAP visualization of scRNAseq cells with boxes colored by cell type (Dark red = high expression; light grey = low expression).

**Figure S3:**
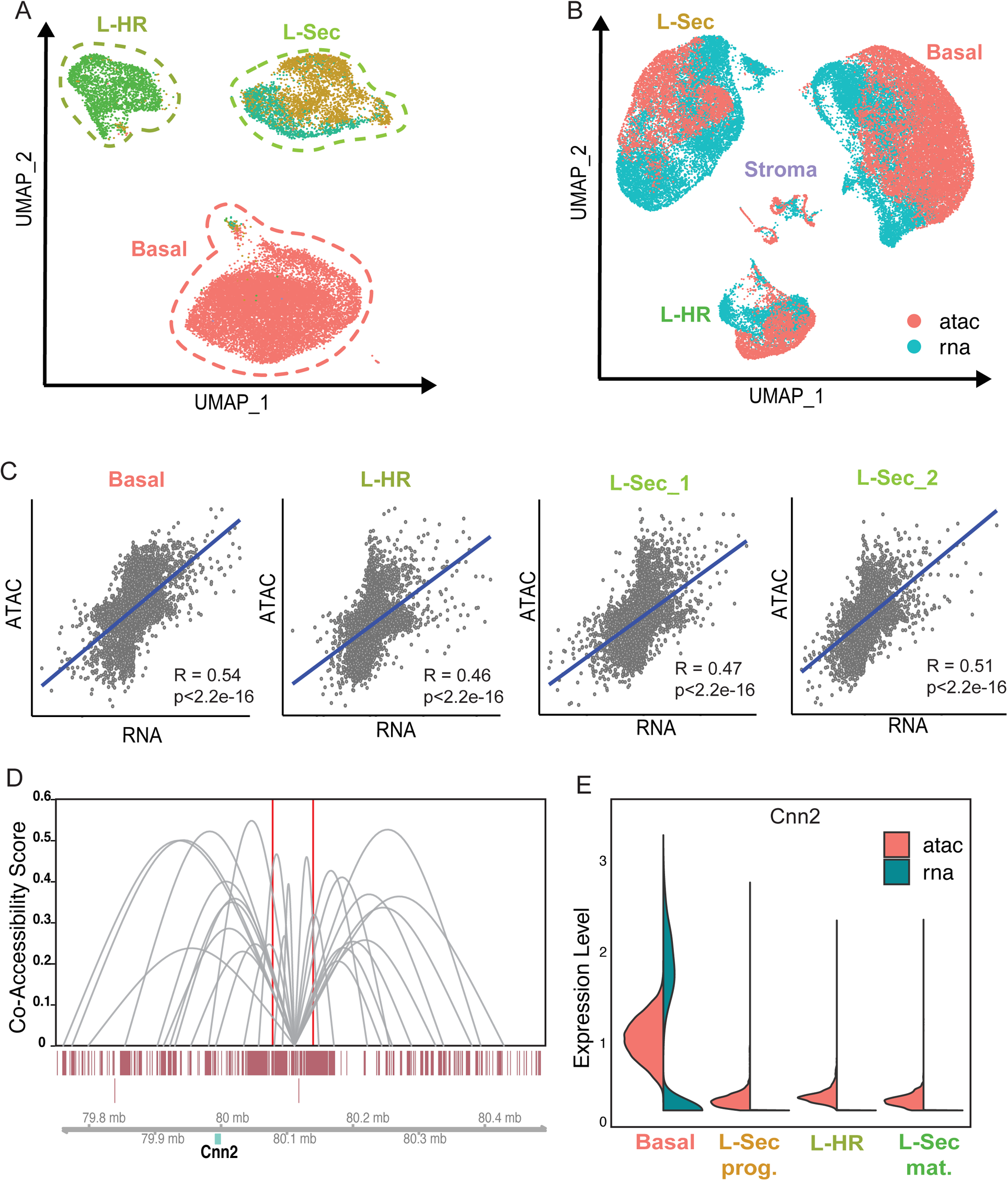
(A) UMAP representation of scATACseq data, with cells colored by predicted cell type label transferred from scRNAseq data clustering. (B) Coembedded UMAP representation of both scATACseq cells and scRNAseq cells, with colors corresponding to the data type of origin. (C) Correlation of gene activity matrix from scATACseq cells and gene expression in cells from scRNAseq, split by cell type. (D) Cicero connection data at enhancer region in proximity to *Cnn2* locus. Connections within Basal cells are shown, with a minimum co-access score of 0.2 visualized. (E) Violin plot of *Cnn2* expression in the coembeded analysis, split by technology type.

**Figure S4:**
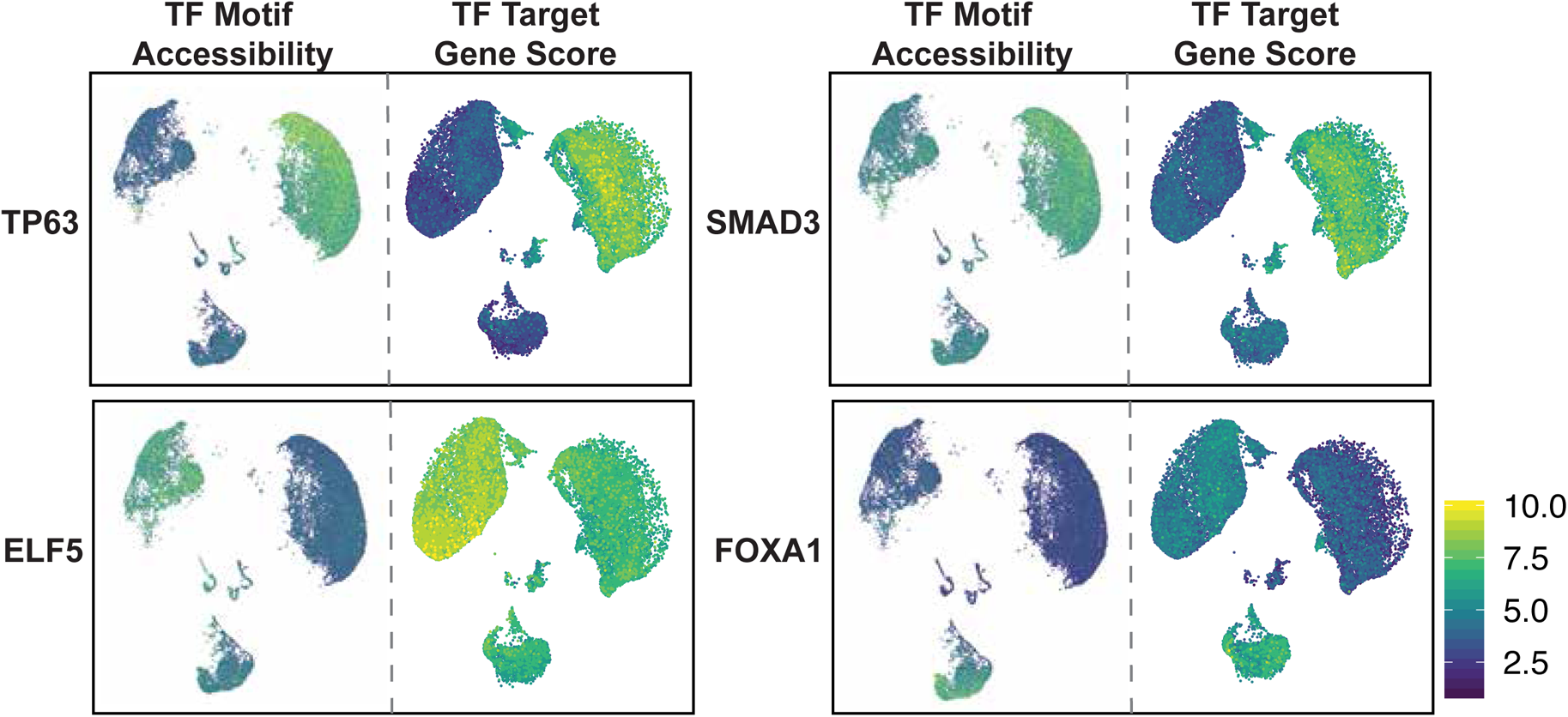
Faceted UMAP visualization of coembedded analysis, with scATACseq cells on the left, and scRNAseq on the right. scATACseq data is colored by z-scored deviations of TF motif accessibility, and scRNAseq data is colored by gene scoring of downstream targets of the TF signaling as annotated through GO Terms. Yellow corresponds to high values, and dark blue to low.

